# Optimizing touchscreen measures of rodent cognition by eliminating image bias

**DOI:** 10.1101/2021.04.05.438342

**Authors:** James A. Belarde, Claire W. Chen, Elizabeth Rafikian, Mu Yang, Carol M. Troy

**Affiliations:** Department of Neuroscience, Columbia University Vagelos College of Physicians and Surgeons, New York, NY 10032, USA; Department of Pathology & Cell Biology, Columbia University Vagelos College of Physicians and Surgeons, New York, NY 10032, USA; Institute for Genomic Medicine, Columbia University Vagelos College of Physicians and Surgeons, New York, NY 10032, USA; Department of Neurology, Columbia University Vagelos College of Physicians and Surgeons, New York, NY 10032, USA; The Taub Institute for Research on Alzheimer’s Disease and the Aging Brain, Columbia University Vagelos College of Physicians and Surgeons, New York, NY 10032, USA

## Abstract

For the last twenty years, the Bussey-Saksida touchscreen-based operant conditioning platform has evolved in close parallel alongside the Cambridge Neuropsychological Test Automated Battery (CANTAB) to produce batteries of tests for studying complex cognitive functions in rodents that are increasingly analogous to human diagnostic tests and greatly narrow the translational gap in cognition research. Naturally, with this increasing usefulness comes increasing use, particularly by non-experts. This necessitates a greater understanding of, and a better controlling for, confounding factors that may limit the system’s ability to optimally detect cognitive deficits when used as a widely accessible and commercially available standardized task. In the present study, we show a strong image preference bias in a standard pairwise discrimination task with a widely used spider-plane image pairing in a putative animal model for intellectual disability. This bias greatly influenced the performance of our experimental mice, significantly affecting the length of time it took mice to complete the task, their progress over time, and several accessory measures usefully recorded by the Bussey-Saksida touchscreen system. We further show that this bias can be corrected by using more similar image pairings without sacrificing the animal’s ability to learn to distinguish the stimuli. This approach eliminated all significant stimuli specific differences seen with the spider-plane pairing. We then analyzed the pixel composition of the various stimuli to suggest that the bias is due to a difference in image brightness. These findings highlight the importance of carefully modulating paired touchscreen stimuli to ensure equivalence prior to learning and the need for more studies of visual perception in mice, particularly as it relates to their performance in cognitive assays.

## 1 INTRODUCTION

Cognitive processes are an assortment of distinct but interrelated brain functions vital to learning about and making sense of the world around us, as well as exploiting that knowledge in goal-directed behavior. Often seen as the faculties that make humans more advanced than other species, these include memory, attention, communication, reasoning ability, and problem solving, as well as others. These important cognitive capabilities are also susceptible to dysfunction from a wide variety of neurological insults including neurodevelopmental disorders, trauma, psychiatric illness, neurovascular damage, infection-induced delirium, autoimmune disease, dementia, and Alzheimer’s Disease. As such, cognition studies are vital for translational neuroscience research.

Rodent models are useful preclinical tools to study phenotypes relevant to understanding human diseases such as those listed above. However, the diversity of cognitive impairment and the notable species differences in all levels of cognitive processing pose a unique challenge for reliably evaluating normal and impaired cognitive functions in animal models ^1^. Classic learning and memory paradigms, such as fear conditioning and water maze, while having contributed enormously to our basic understanding of neural mechanisms of learning and memory, have the disadvantage of relying on the use of aversive stimuli ^2,3^. These tasks are also limited when it comes to studying complex cognitive functions and have limitations to their translational value ^4^. Low-stress tests such as novel object recognition suffer from inconsistency in scoring criteria and the difficulty in controlling the amount of “learning” (i.e. time spent with each object) during the learning session ^5^.

The Bussey-Saksida touchscreen system was developed to overcome these limitations and increase the translational value of rodent cognition research. In the touchscreen-based operant conditioning platform, rats or mice are trained to interact with visual stimuli projected on a touch-sensitive screen in various tasks, and correct responses are rewarded with palatable liquid or food. The origin of the touchscreen system traces back to the invention of the *Cambridge Neuropsychological Test Automated Battery* (CANTAB) by Trevor Robbins and colleagues in the 1980s. For over 30 years, CANTAB has been used worldwide to evaluate cognitive impairments in human neurodevelopmental, neurodegenerative, and neuropsychiatric disorders^6^. In collaboration with Trevor Robbins, Tim Bussey and colleagues then developed the first touchscreen system specifically for rats ^7^, which was soon after adapted for mice ^8^. Over the last two decades, the Bussey-Saksida touchscreen system has evolved in close parallel with CANTAB, creating a battery of tests to study complex cognitive functions in ways that are technically and conceptually similar to human versions of these tasks, making it more possible than ever to bridge the translational divide in cognition research ^9,10^.

The benefits of the touchscreen system for testing cognitive function are numerous and have been discussed elsewhere ^9,10^. One major appeal is that the system appears to be “plug-and-play” ready: with standardized equipment, software, and individual task programs available for purchase, as well as users’ manuals and published protocols available for download, one is inclined to assume that following all the steps is both critical and sufficient for obtaining valid experimental results. The simplicity of this user-friendly platform has led to a surge in touchscreen-based cognitive studies in rodents across many disease models including autism ^11,12^, Alzheimer’s Disease ^13,14^, schizophrenia ^15,16^, and Huntington’s Disease ^17^. With the increasing popularity and accessibility of this system, it is important to ensure that possible confounding factors are identified and, where possible, corrected to avoid inaccurate findings. This is especially important information for investigators like the authors of this present study: those that are new to or infrequent users of the touchscreen system and lack the experience of experts in the field.

One aspect of the touchscreen “package” that is usually taken for granted by casual users is the pairs of images used. One easily assumes that each pair of images given as examples in the original protocols ^18,19^ must be equally preferred by the animals in the unrewarded condition before any association learning. While we are not aware of any systematic analysis on image preference/bias in commonly inbred strains, personal communication with several touchscreen experts indicate that the image bias issue is far from a rarity, and seems to be influenced by factors such as images, mouse strain ^20^ and testing location/facility. In the original mouse study ^8^, it was stated that the stimuli (white patterns on black background) were selected to be “approximately equiluminescent”, and to be approximately equal in size (4×4 cm). Even so, to our knowledge, there has been no systematic analysis of often-used image pairings to ensure they are entirely, rather than approximately, equivalent in visual properties like luminescence. This is an important thing to ensure as a salient bias could introduce significant noise into the data and mask subtle but significant effects, resulting in either inaccurate or inconclusive assessments and thereby compromising reliability, sensitivity, and resolution of the touchscreen tasks.

In this study, we used the standard Bussey-Saksida visual discrimination task to test a mouse model with homozygous *cradd* deletion (on the C57BL/6L background), a putative model of a human disorder characterized by intellectual disability ^21^. As non-experts in the field of mouse behavioral cognition, we were drawn to this method for all the reasons discussed above: ease of use, reliance on appetitive rather than aversive stimuli, and the apparent “plug-and-play” approach of a standardized model. We report here the incidental finding of an image bias: subject mice learned the task significantly faster when the spider of the spider-plane pair is rewarded, regardless of sex or genotype, even with the images optimally counter-balanced. It is important to note that the *cradd* KO animals show no signs of decreased visual capability. An analysis suggested the same bias in an identical test in three other mouse lines tested at our facility, suggesting this bias is not limited to the *cradd* KO experimental animals. This surprised us, since we had chosen this particular image pairing included in the software given the large number of published studies that utilized the spider-plane pair for visual discrimination and reversal ^11,12,19^, or related paradigms including transitive inference ^22^ and paired-associate learning (PAL) ^23-26^. We further demonstrate that this bias affects multiple measures across both the acquisition and reversal training phases of the pairwise discrimination task. We then show that the image bias can be eliminated by using identical images constituted of black/white stripes turned to different angles (X and X + 180°), as have been used in several previous studies ^19,27,28^, without significantly affecting overall performance. Finally, we suggest that the observed image bias could be due to brightness differences as the result of an unintended learned association established in a standard pretraining protocol, highlighting the importance of carefully modulating this part of the experiment as needed.

Our study highlights an important, yet, as far as we’re able to ascertain, under-addressed issue of image bias in visually based touchscreen tests of rodent cognition. These findings call for more comprehensive, definitive studies by touchscreen experts and rodent behaviorists to identify and eliminate bias in standardized protocols intended for more general use by casual touchscreen users. Our data also highlight the importance of studying visual perception in mice and the ethological relevance of images presented to them, especially those stimuli that are not composed of simple geometric shapes, in order to more accurately interpret visual cognitive behaviors in mouse models.

## 2 METHODS

### 2.1 Animals

*Cradd*^-/-^ mice on a C57BL/6J background were obtained from the Jackson Laboratories (Bar Harbor, ME) with permission from Prof. Tak Mak (University of Toronto). Heterozygote *cradd*^+/-^ mice were bred to generate homozygous *cradd*^-/-^(KO) experimental animals, as well as wildtype (WT) littermate controls. Prior to experiment onset, all experimental mice were maintained in group housing on a 12-hour light/dark cycle with ad libitum access to food and water. All behavioral tests were run during the light cycle between 11am and 3pm every day and adult mice (age 5-8 months) of both sexes were used, including both WTs and KOs. All experiments and protocols were approved and monitored by the Institutional Animal Care and Use Committee (IACUC) of Columbia University.

### 2.2 Touchscreen pairwise discrimination and reversal

Pairwise visual discrimination was tested in the automated Bussey-Saksida touchscreen apparatus for mice (Campden Instruments Ltd/Lafayette Instruments, Lafayette, IL, USA), using a procedure modified based on original methods described previously^4,8,22,29-32^. The reinforcer was 20µl of a palatable liquid nutritional supplement (Strawberry Ensure Plus, Abbott, IL, USA) diluted to 50% with water. Each session is conducted under overhead lighting (∼60 lux). A standard tone cue was used to signal the delivery of the reinforcer during pre-training and acquisition. Prior to pre-training, subject mice were weighed, and placed on a restricted diet of 2-4g of rodent chow per mouse per day, to induce 10-15% weight loss. Body weight was carefully monitored throughout the experiment to ensure that a minimum of 85% of free-feeding body weight was maintained for each mouse. Mice were maintained in the group housing they had occupied since weaning throughout the experiment. All equipment used was thoroughly cleaned, disinfected, deodorized, and then wiped down with water in between each mouse. Mice were trained 5 days a week with one 1-hour session per day.

We used a 5-stage pre-training regimen based on pioneering methods used previously ^4,12,18,22,29,30^. In order, these stages are habituation, initial touch, must touch, must initiate, and punish incorrect. For habituation, the mice are introduced to the chamber and liquid reinforcer for 40 minutes on two consecutive days. No images are displayed in this first stage. During initial touch, an image is presented on one of the two screens. If the image is not touched within 30 seconds, the touchscreen panel is turned off and 20µl of liquid reinforcer is delivered to the reward tray, while touching the image turns off the touchscreen and is rewarded with 3x the amount of reinforcer, signaling a successful trial completion. To advance to the next stage, mice must complete 30 such trials within a 1-hour testing period. In the must touch stage, an image is displayed on one of the two screens and remains on until touched, at which time the liquid reward (20µl) is delivered. Mice had to complete 30 such trials within the 1-hour session for two consecutive days to advance to the next stage. The must initiate stage required the mouse to nose-poke the reward magazine to initiate the next trial, at which point an image is displayed on one of the two screens until it is touched, and the liquid reward delivered. Again, mice must complete 30 trials successfully in a 1-hour period for two consecutive days to advance. In the final pretraining stage, punish incorrect, touching the blank screen with no image when an image is present on the opposite screen is punished with a 5-second timeout. To pass this stage, mice must complete 30 trials in the 1-hour time limit with 80% accuracy or better for two consecutive days. Only mice that completed all five of these stages were advanced to the pairwise visual discrimination task. Images used in pre-training were randomized selections from a pre-training image library that did not include any images used in the subsequent discrimination task. One WT female was dropped from the spider-plane image pairing experiment after failing to complete the initial touch stage within the 5-day cutoff. All other mice in both experiments successfully completed pre-training.

For pairwise discrimination, subjects were trained to discriminate between two novel images, a spider and an airplane, presented in a spatially pseudo-randomized manner in the two windows of the touchscreen. Each 1-hour session consisted of 30 trials separated by 15-second intertrial intervals (ITI). Designation of the correct and incorrect images was counterbalanced across mice within each genotype. Correct responses were rewarded. Each incorrect response was followed by a correction trial in which the images were presented in an identical manner to the previous trial, until a correct response was made. Criterion was completing all 30 trials at an accuracy of 80% or higher over 2 consecutive days.

Reversal training began approximately 3 days after the last acquisition day. In the reversal task, the designation of correct and incorrect images in the acquisition phase was reversed. As in acquisition, the criterion was an average of ≥ 80% correct responses on 2 consecutive days. For acquisition, a 20-day cutoff was applied, while a 40-day cutoff was used for the reversal stage. For the purposes of data analysis, animals that failed to reach criterion in 20 days during the acquisition phase were dropped from the study and not included in analysis; no mice were dropped after acquisition in the spider-plane image pairing experiment, while 2 WT male mice were dropped in the mirrored diagonals experiment. Days to reach criterion, percentage of mice reaching criterion, number of errors, correction errors, correct trials, total trials, and inactive touches (interactions with touchscreen during ITIs when no stimulus is present) were compared between groups designated by their correct image assignments for both the acquisition and reversal phases.

### 2.3 Image pairs used

The following are explanations of the three image pairings used in our study **(Fig. 1)**. For all pairings, the image designated as the correct (or rewarded) image was counterbalanced across all animals in the acquisition and reversal phases. The presentation location for the two images was pseudorandomly selected between trials, allowing each stimulus to be presented equally in both response locations across every testing session. Two experiments were conducted using two separate cohorts of mice from the same *cradd* KO line, designated Cohort 1 and Cohort 2 below.

**Figure 1:**
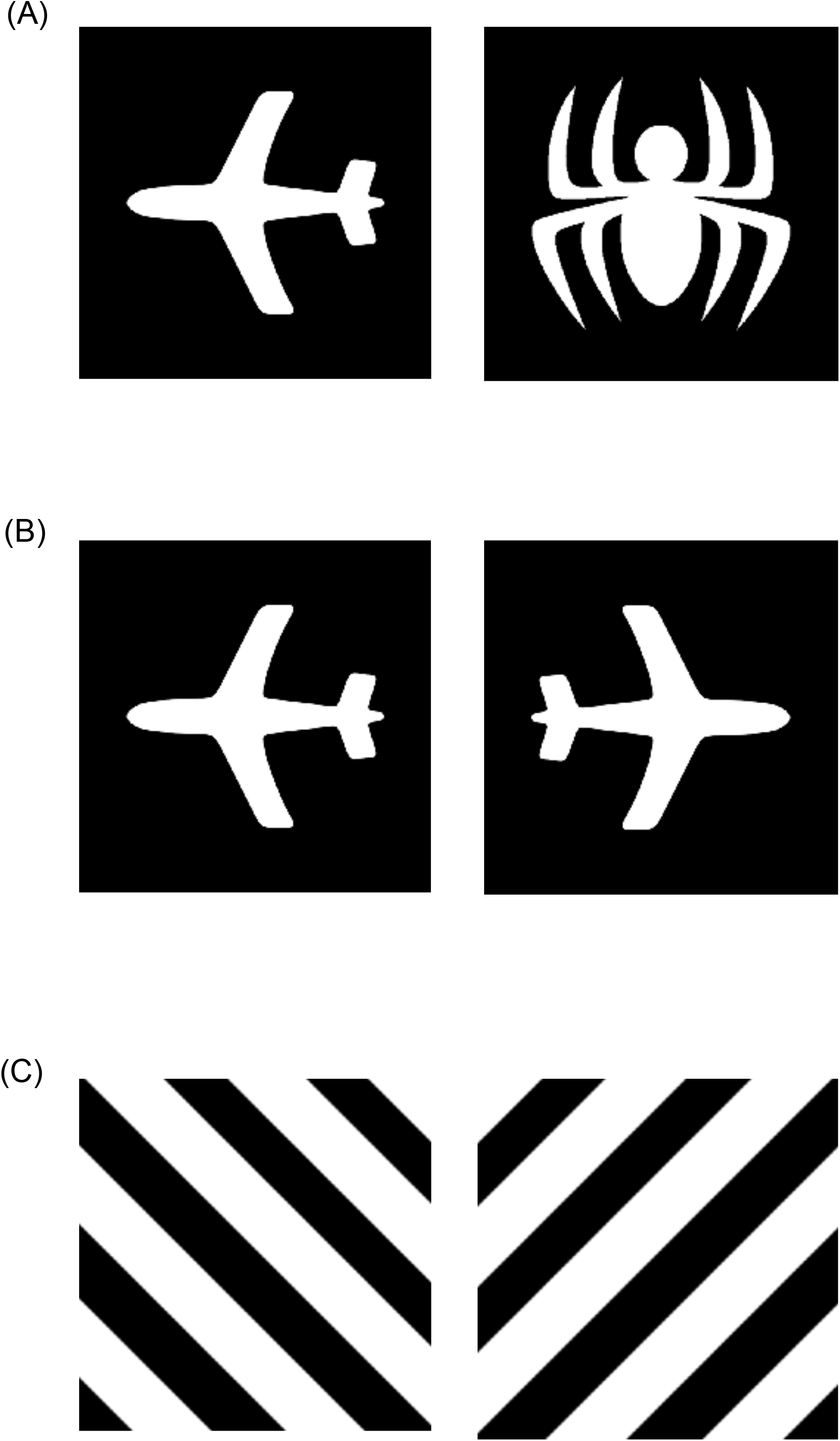
Three image pairings were used in this study to assess image bias. (A) One of the two most commonly used image pairings in touchscreen publications, the spider-plane image pairing is included in the Bussey-Saksida Touchscreen system software. (B) Rotating the standard plane image from A by 180°, we utilized a pairing of the plane image and its reflection. (C) An image pairing using a repeating diagonal bar pattern and its 180°-rotated reflection, as described in Phillips et al., 2017^28^.

#### Spider and plane

Along with marble-fan, this is one of the two most commonly used image pair standards in touchscreen experiment protocols, featuring a simple solid white outline of either a spider or a plane (Fig. 1A). This pairing is featured as an option in the software accompanying the Bussey-Saksida Touchscreen system and was used for the Cohort 1 experiment.

#### Plane and mirrored plane (Plane+180o)

For this image pairing, we used the standard plane image as in the spider-plane pairing, as well as that same plane image rotated 180° such that it was facing the opposite direction as the original **(Fig. 1B)**. We chose the plane image for this because its decreased symmetry (as compared to the spider) suggested a greater chance the mice would still be able to distinguish between the reflected images. This pairing was initially used for the Cohort 2 experiment. After 10 days of training showed no improvement in the animals’ performances, they were given a 2-day break and switched to mirrored-diagonal stripes as described below.

#### Mirrored-diagonal stripes

After Cohort 2 failed to effectively distinguish stimuli in the plane-mirrored plane pairing, we sought alternative images with equal similarity but different enough to allow for discrimination. In studies with head-fixed mice, identical stripes positioned as X and X+45° or 90° were effective in eliciting significantly different responses from mouse V1 cortex ^33^, suggesting that the differences between the two images are biologically detectable. This is further supported by a study in which a cross and a cross rotated 45 degrees were used to study visual discrimination ^34^, and others which have successfully utilized striped stimuli for this task ^28,35^.

For our experiment, we used the diagonal bands of black and white stripes as described in Phillips et al. **(Fig. 1C)**. These images were designated as either left diagonal (LD; diagonal bars ran from top left corner to bottom right corner) or right diagonal (RD; diagonal bars ran from top right corner to bottom left corner).

### 2.4 Statistical Analysis

All data other than Kaplan-Meier curves are presented as means ± standard error of the mean (SEM). Scatterplots included in some graphs represent datapoints from individual subjects that were used to calculate the indicated averages. Kaplan-Meier curves are used to visualize “time-to-event” data, here indicating the cumulative percentage of mice that have reached criterion (see **Methods 2.2**) and completed the touchscreen task by a given experiment day. Kaplan-Meier curves were statistically analyzed using a log-rank test. All other data was analyzed using a Welch’s t-test for unequal variances.

## 3 RESULTS

### 3.1 Experimental animals show a significant image bias when standard spider-plane stimulus pairing used

Our study began as an attempt to ascertain cognitive defects in our *cradd* KO mouse model using a touchscreen-based pairwise discrimination protocol. We started with a common spider-plane image pairing as included in the software for the Bussey-Saksida Touchscreen system **(Fig. 1A)**. In this first study, we saw no differences in either the acquisition or reversal phase of the behavioral task when comparing WT and KO genotypes (data not shown). However, the experimenters noted that, regardless of genotype, mice appeared to reach criterion faster in both phases when the rewarded image was spider. To quantify this observation, mice were regrouped into plane-correct (Plane+; N = 13 for acquisition: 4 WT males, 3 KO males, 2 WT females, 4 KO females; N = 11 for reversal: 3 WT males, 2 KO males, 2 WT females, 4 KO females) and spider-correct (Spider+; N = 11 for acquisition: 3 WT males, 2 KO males, 2 WT females, 4 KO females; N = 13 for reversal: 4 WT males, 3 KO males, 2 WT females, 4 KO females) groups, and the data were reanalyzed. The Spider+ group reached criterion significantly faster than the Plane+ group for both acquisition (**Fig. 2A**; Days to criterion: Spider+: 7.91±0.58; Plane+: 10.08±0.57; p = 0.02) and reversal (**Fig. 2B**; Spider+: 9.00 ± 0.46; Plane+: 12.45 ± 0.75; p = 0.002). It is important to note that, due to the nature of the task, the Spider+ group in the acquisition phase of the experiment becomes the Plane+ group in the reversal phase and vice versa, strongly suggesting this effect is driven by the specific image rather than any factor inherent in our grouping of the mice.

**Figure 2:**
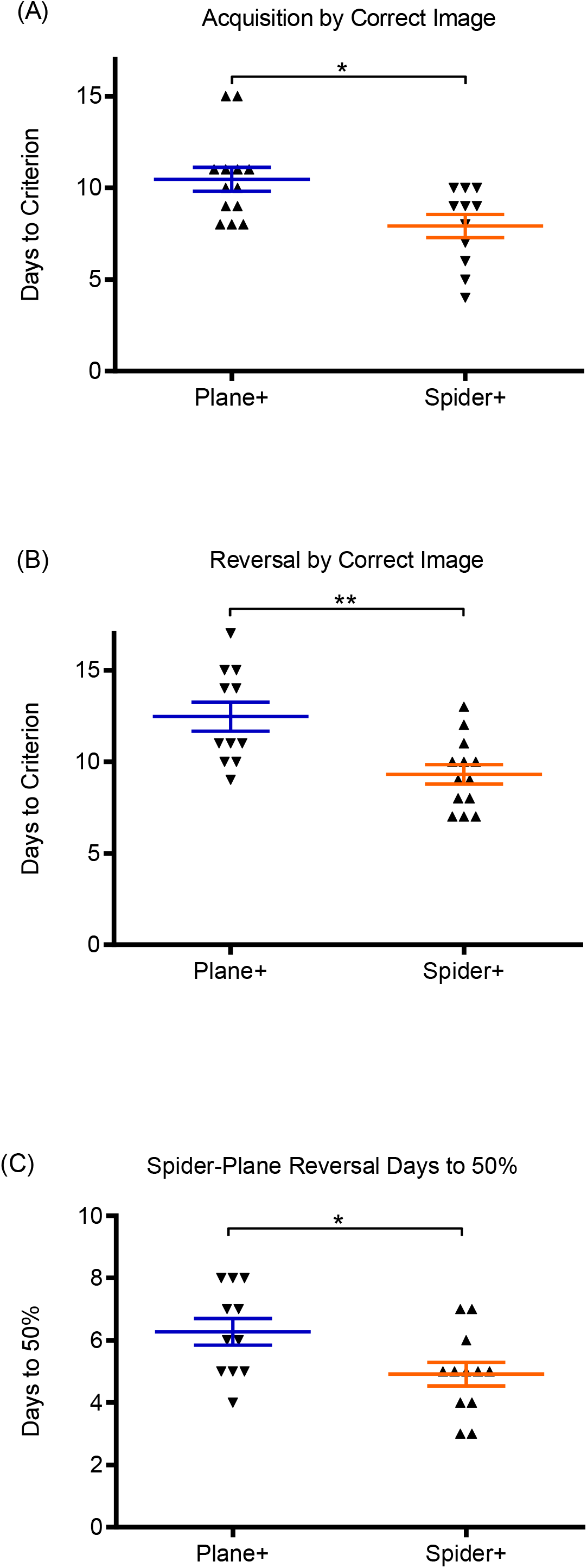
Spider-correct (Spider+) groups reach criterion significantly faster than plane-correct (Plane+) groups in both (A) acquisition and (8) reversal stages of a pairwise discrimination task. (C) The Spider+ group also returns to a baseline chance performance of 50% significantly faster than the Plane+ group during reversal phase. Data are presented as mean ± SEM, with scatterplots representing the individual scores used to obtain those means. Upward arrows represent one consistent group of mice throughout the task, while downward arrows represent the other group. *p < 0.05, **p < 0.01

According to the literature, another important parameter in the reversal phase of the pairwise discrimination task is days to 50% (DT50), when the mice return to a baseline of chance performance and touch the correct image 50% of the time. This is a measure of the length of time it takes for a subject to unlearn a previously acquired rule and adopt a new approach when the rules are switched. The Spider+ group had a significantly lower DT50 value than the Plane+ group (**Fig. 2C**; Spider+: 4.92±0.36; Plane+: 6.27±0.41; p = 0.03), indicating that mice in the Spider+ group spent less time perseverating on the previously acquired rule when the correct image had been switched.

### 3.2 Image bias creates a significant difference in how the two groups perform over time

Given that pairwise discrimination experiments are conducted over several weeks, Kaplan-Meier curves and averaged learning curves are useful ways to visualize the data more temporally. In both acquisition and reversal, there is a pronounced leftward shift in the Kaplan-Meier curve for the Spider+ groups as compared to Plane+ ones (**Fig. 3A** and **3B**). Running a log-rank statistical analysis to compare these curves showed that this difference was significant in both phases of the task (acquisition p = 0.03; reversal p < 0.01). This significant shift shows a clear bias toward the spider image from early in the experiment.

**Figure 3:**
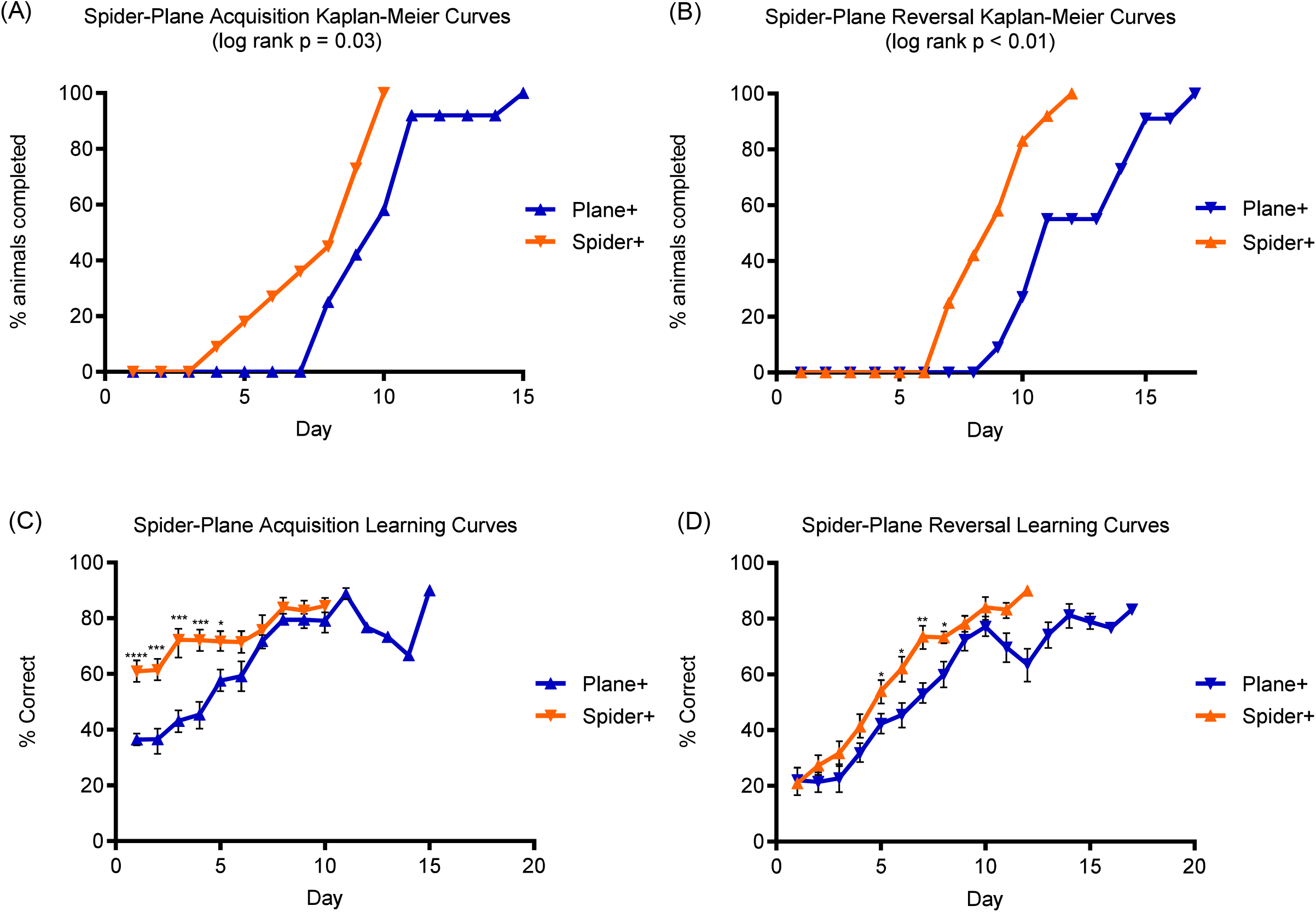
Spider+ groups show significantly faster progress in the task over time as compared to Plane+ groups. (A) and (B) Kaplan-Meier curves were obtained by plotting the cumulative percentage of mice in either group that had completed the task by a given day. There is a significant leftward shift of the Spider+ group’s Kaplan-Meier curve as compared to the Plane+ group in both acquisition (A) and reversal (B). In (C) and (D), averaged learning curves were obtained by averaging the % correct score across all mice in each group for the indicated day. The Spider+ group performed significantly better than the Plane+ group on multiple days throughout in both acquisition (C) and reversal (D). For Kaplan-Meier curves, indicated p-values were obtained using a log-rank analysis. Averaged learning curve data are presented as mean ± SEM, *p < 0.05, **p < 0.01, ***p < 0.001, ****p < 0.0001

The averaged learning curves highlight this further **(Fig. 3C** and **3D)**. These curves are obtained by averaging the percent correct of both groups for each day. In the acquisition phase, the bias for the spider image is apparent as early as day 1. Under normal conditions, one would expect the animals to perform at or near chance (50% correct) when the task is first presented. However, the Spider+ group is already performing at around 60% correct on average on day 1, while the Plane+ group averages under 40% correct. These differences were significant for several days at the start of acquisition (**Fig. 3C**; Day 1: p < 0.0001; Days 2-4: p < 0.001; Day 5: p = 0.018; all other days: p>0.05). In the reversal phase of the task, both groups start at around 20% correct on day 1, showing that both are perseverating on the previously correct image regardless of which that was. However, the Spider+ group then proceeds to outperform the Plane+ group on all following days of reversal, differences that again reached significance at several different timepoints as Spider+ group’s progress began to outpace that of Plane+ (**Fig. 3D**; Day 5: p = 0.048; Day 6: p = 0.025; Day 7: p < 0.01; Day 8: p = 0.019; all other days: p > 0.05).

### 3.3 Spider-correct group outperforms plane-correct group across multiple measures

Several other measures tracked by the touchscreen apparatus also illustrate a choice bias in favor of the spider image. While both Spider+ and Plane+ groups completed the same number of trials overall in acquisition (**Fig. 4A**; Spider+: 235.73±17.18; Plane+: 274.62±23.93; p = 0.20) and reversal (**Fig. 4B**; Spider+: 228.38±18.87; Plane+: 278.73±22.09; p = 0.11), the Spider+ group required significantly fewer correction trials (**Fig. 4C**; Spider+: 121.91±19.66; Plane+: 228.08±14.03; p = 0.00047) and made significantly fewer errors (**Fig. 4E**; Spider+: 66.36±8.58; Plane+: 99.85±8.31; p = 0.01) than the Plane+ group in the acquisition phase. During reversal, these differences failed to reach significance for both number of correction trials (**Fig. 4D**; Spider+: 317.31±27.18; Plane+: 361.27±21.77; p = 0.23) and number of errors (**Fig. 4F**; Spider+: 100.00±11.51; Plane+: 117.55±9.37; p = 0.26), possibly due the greater difficulty of this stage of the task.

**Figure 4:**
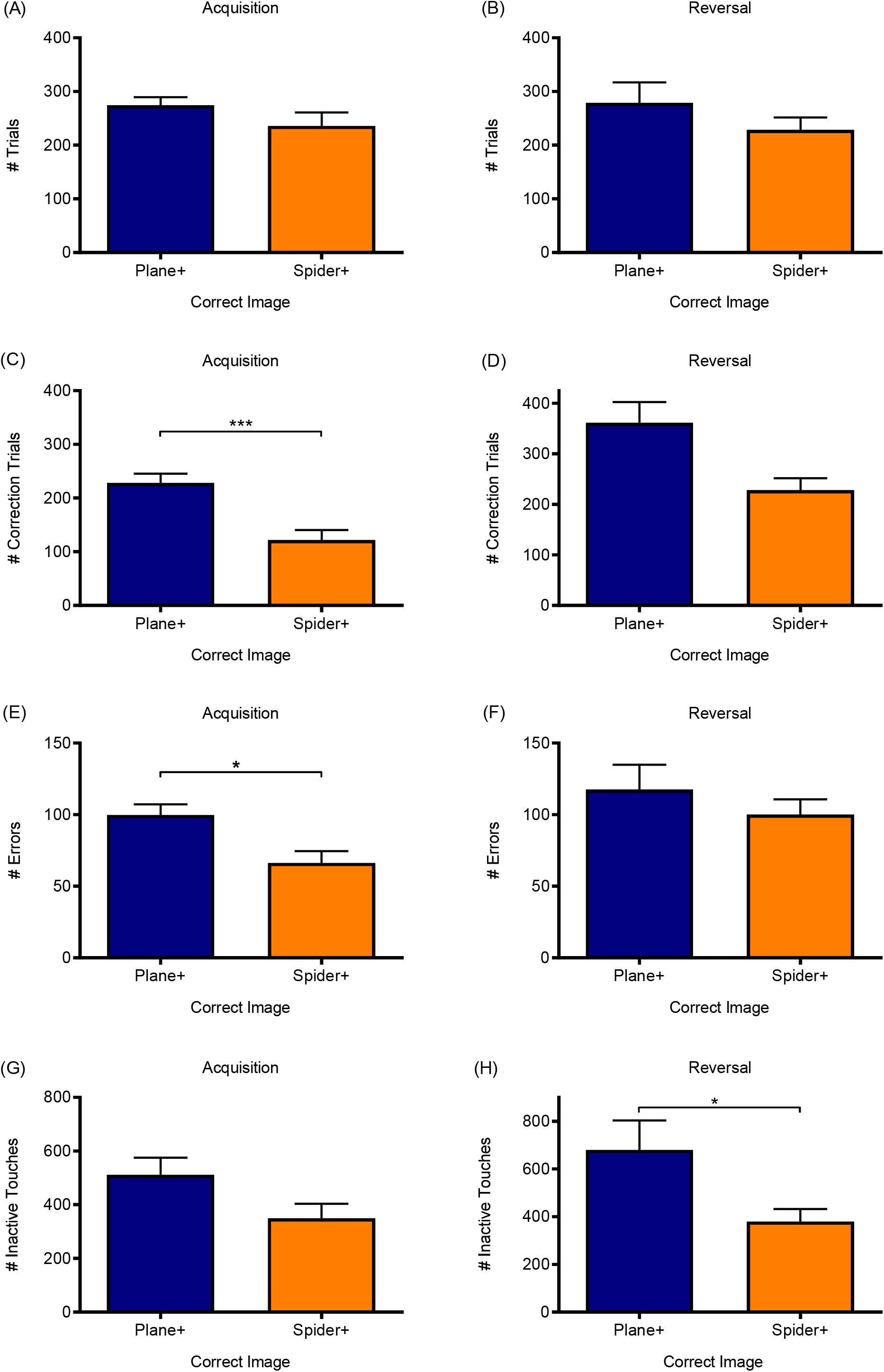
Spider+ groups (orange bars) perform better than Plane+ groups (blue bars) by several accessory measures examined. On average, Plane+ and Spider+ groups completed an equivalent number of total summed trials in both acquisition (A) and reversal (8). Despite this, the Plane+ group required significantly more correction trials (C) and made a significantly greater number of errors (E) overall than the associated Spider+ group in acquisition. These differences failed to reach significance in the reversal stage for both total correction trials (D) and total errors made (F). The Spider+ group also makes fewer inactive touches (interactions with the touchscreen when no image is displayed and thus no choice being made) than the corresponding Plane+ group. While this observation just narrowly failed to reach significance in acquisition (G; p = 0.07), the difference was significant in reversal (H). Data are presented as mean± SEM, *p < 0.05, ***p < 0.001

Interestingly, the Spider+ groups also made fewer inactive touches on average than their corresponding Plane+ groups, a difference that narrowly failed to reach statistical significance in acquisition (**Fig. 4G**; Spider+: 349.91±51.29; Plane+: 511.92±68.33; p = 0.07), but was significant in the reversal phase (**Fig. 4H**; Spider+: 380.15±55.78; Plane+: 679.73±118.94; p < 0.05). This measure tracks the number of times the mouse touches the screen during the intertrial intervals, when no images are presented.

### 3.4 Using plane-mirrored plane stimulus pairing fails to elicit image discrimination

After seeing this unexpected finding of the above-mentioned image bias in our mouse model, we decided to use a task-naïve group of mice from the same line to explore the possibility of eliminating this bias. To do so, we decided to modify the standard images such that they would be as similar as possible while still being distinguishable. For this, we chose to use the plane and its reflection as the two stimuli **(Fig. 1B)**. As can be seen from the average learning curves **(Fig. 5)**, using these mirrored plane images as the stimulus pair eliminated the bias and both groups (N = 9/group) started near the expected 50% chance performance, with no significant differences between groups on any day (p>0.05 for all 10 days). However, the lack of bias in this case is purely secondary to the fact that the animals failed to discriminate the strikingly similar images at all, failing to reach criterion or show any appreciable improvement in their learning curves after ten days of testing.

**Figure 5:**
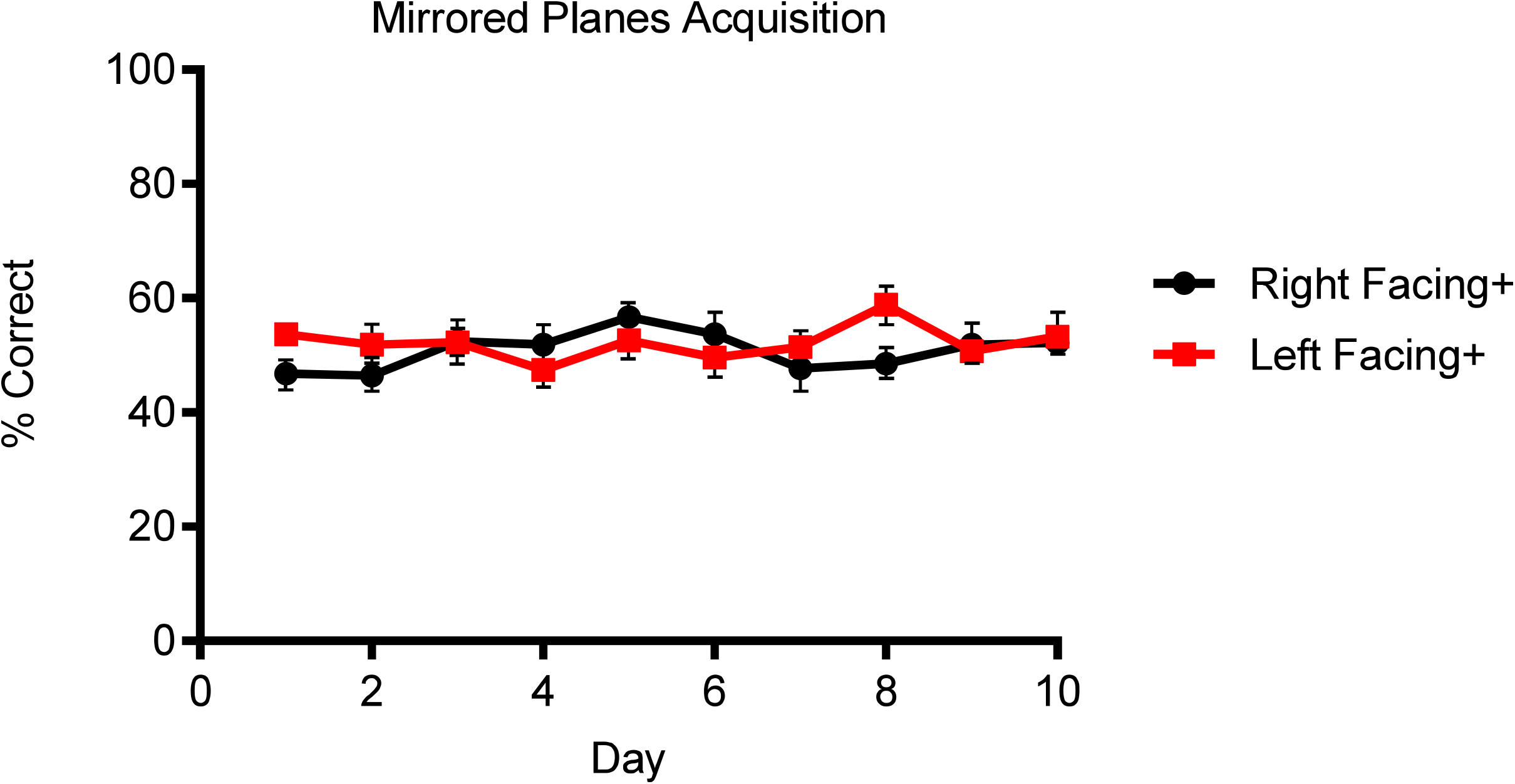
Use of mirrored plane image pairing eliminates any evident image bias. This is seen in the near complete overlap of the two groups’ curves, with no significant differences on any day. However, failure to advance above chance performance (50%) after ten days suggests this pairing prohibits successful completion of the discrimination task. Data are presented as mean ± SEM

### 3.5 Mirrored diagonal bar pattern stimuli eliminate bias in multiple measures while still allowing mice to effectively complete the task

After realizing that the mirrored plane stimuli were not sufficiently distinct for our mice to learn the task, we sought an image pairing that would still be similar enough to avoid bias but provide enough salient differences to allow learning. This led us to obtain a set of mirrored diagonal bar patterns to use as the stimulus pair **(Fig. 1C)**, identified as either left-diagonal or right-diagonal depending on the direction in which the diagonal bars ran from the bottom of the image to the top. Given the image’s uniformity and multiple points of contrast, we predicted the mice would be able to distinguish the two reflected patterns despite the high degree of similarity between the two. And indeed, unlike with the mirrored-plane image pairing, the animals were able to learn to distinguish between the two diagonal stripes images to receive the liquid diet reinforcer. Most importantly, after being separated into left-diagonal correct (LD+; N = 7 for acquisition: 3 WT males, 3 KO males, 1 het female; N = 9 for reversal: 5 WT males, 3 KO males, 1 KO female) and right-diagonal correct (RD+; N = 9 for acquisition: 5 WT males, 3 KO males, 1 KO female; N = 7 for reversal: 3 WT males, 3 KO males, 1 het female) groups, there was no difference in DTC between the groups in either the acquisition (**Fig. 6A**; LD+: 7.88±1.21; RD+: 6.33±0.96; p = 0.36) or reversal (**Fig. 6B**; LD+: 15.67±2.13; RD+: 17.88±3.26; p = 0.59) stage of the task. There was also no significant difference in the cognitive flexibility as measured by DT50 (**Fig. 6C**; LD+: 5.11±0.6; RD+: 5.63±0.42; p = 0.51) between the two groups.

**Figure 6:**
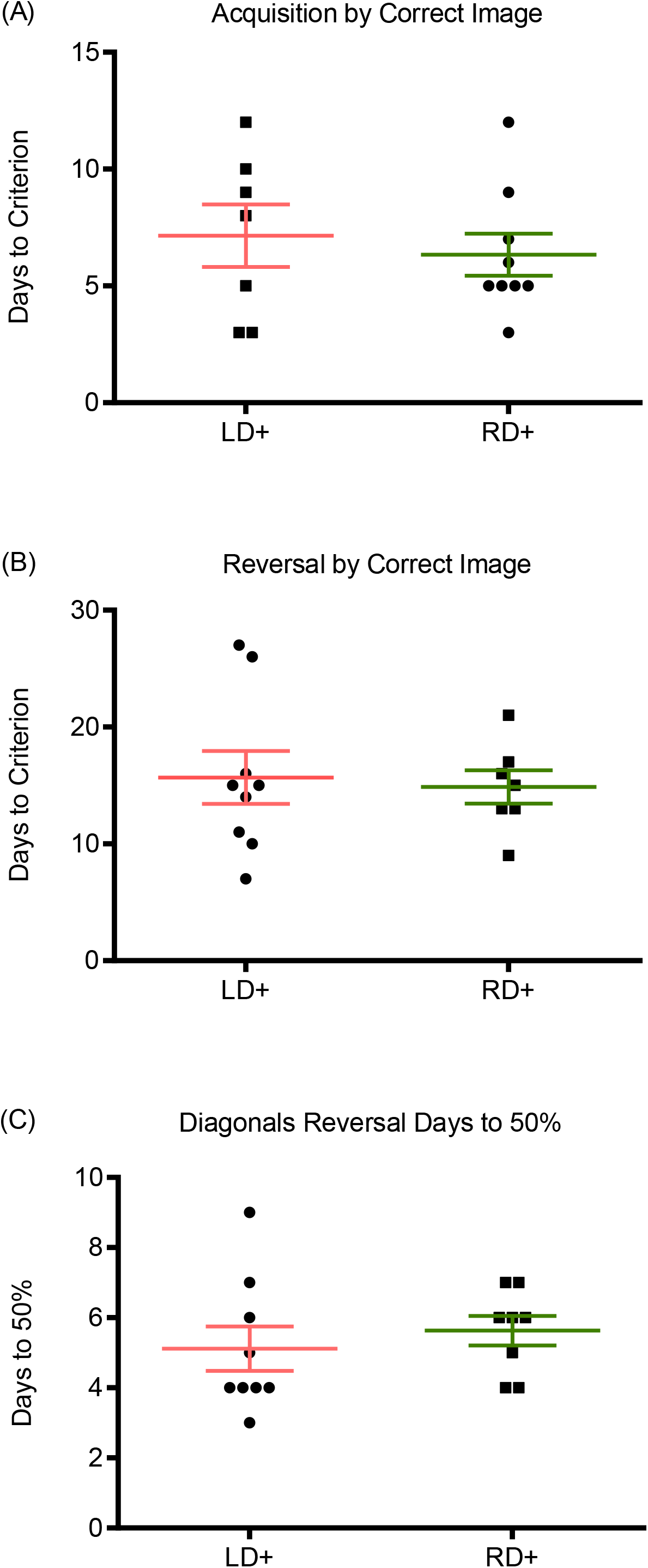
The use of a mirrored diagonal bar pattern image pairing eliminates evident image bias while still allowing for successful discrimination between the two images. Unlike with the spider-plane pairing, there were no significant differences between left-diagonal correct (LD+) and right-diagonal correct (RD+) groups in number of days to reach criterion for either acquisition (A) or reversal (8), nor were there differences in number of days to reach 50% correct in reversal (C). Data are presented as mean± SEM, with scatterplots again representing individual scores. Circles indicate one consistent group of mice throughout the task, while squares represent the other group.

The lack of bias in both phases of the task is also evident from the Kaplan-Meier and learning curves for the mirrored diagonal pairing groups **(Fig. 7)**. The Kaplan-Meier curves for both image groups **(Fig. 7A** and **7B)** overlap without a significant shift in either direction, as indicated by a log-rank analysis (acquisition: p=0.67; reversal p=0.62). Similarly, the learning curves show equivalent progression, with the expected starting points for both acquisition (**Fig. 7C**; groups around 50% correct on day 1, indicating chance performance) and reversal (**Fig. 7D**; under 20% on first day of reversal, indicating a retention of the previous rule after the correct image has now been switched). There was no significant difference between average group performance on any day in either phase (p>0.05 at all timepoints).

**Figure 7:**
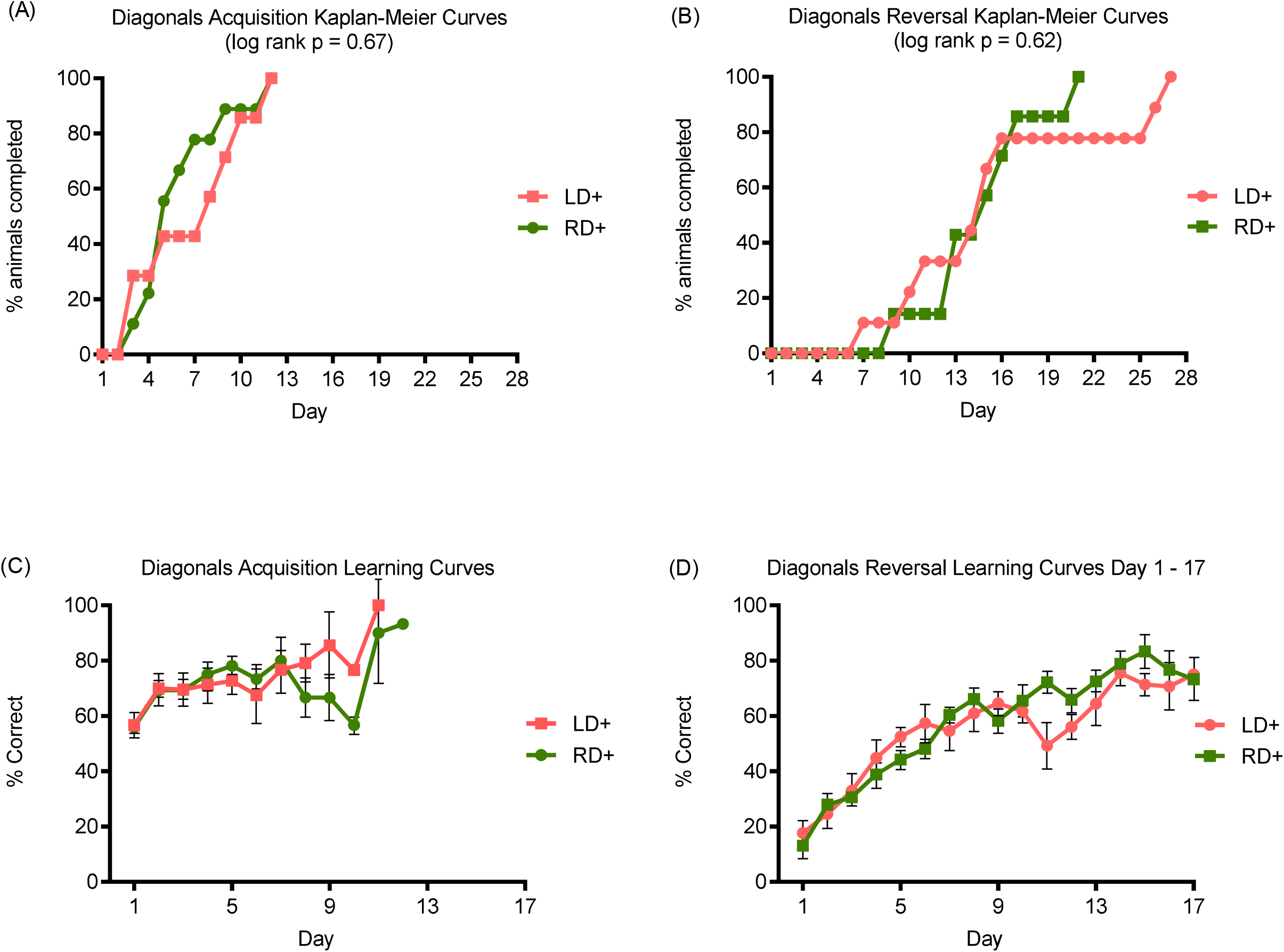
Both the LD+ and RD+ groups show similar rates of progression in the pairwise discrimination task. Kaplan-Meier curves for both acquisition (A) and reversal (B) show appreciable overlap with no significant shift in either curve. Similarly, the averaged learning curves for LD+ and RD+ groups show comparable progress over time, with no significant differences between groups on any day in either acquisition (C) or reversal (D). For Kaplan Meier curves, indicated p-values were obtained using a log-rank analysis. Averaged learning curve data are presented as mean ± SEM

Like in the plane-spider image pairing experiment, both the LD+ and RD+ groups completed similar total numbers of trials in acquisition (**Fig. 8A**; LD+: 218.57±33.26; RD+: 193.33±28.34; p = 0.61) and reversal (**Fig. 8B**; LD+: 394.89±66.16; RD+: 406.43±33.33; p = 0.88). In contrast to the plane-spider experiment, though, there were no significant differences between the diagonal groups in any of the additional measures compared, including correction trials during acquisition (**Fig. 8C**; LD+: 96.71±16.13; RD+: 100.78±28.27; p = 0.90) and reversal (**Fig. 8D**; LD+: 492.89±90.77; RD+: 520.43±37.19; p = 0.78), errors made in acquisition (**Fig. 8E**; LD+: 59.14±9.89; RD+: 56.44±13.93; p = 0.88) and reversal (**Fig. 8F**; LD+: 162.78±26.17; RD+: 178.00±14.36; p = 0.62), and average inactive touches in acquisition (**Fig. 8G**; LD+: 201.14±50.31; RD+: 245.33±51.93; p = 0.52) and reversal (**Fig. 8H**; LD+: 1093.11±253.27; RD+: 778.14±145.72; p = 0.31). These data suggest that both groups perform equally well across the experiment with no confounding bias.

**Figure 8:**
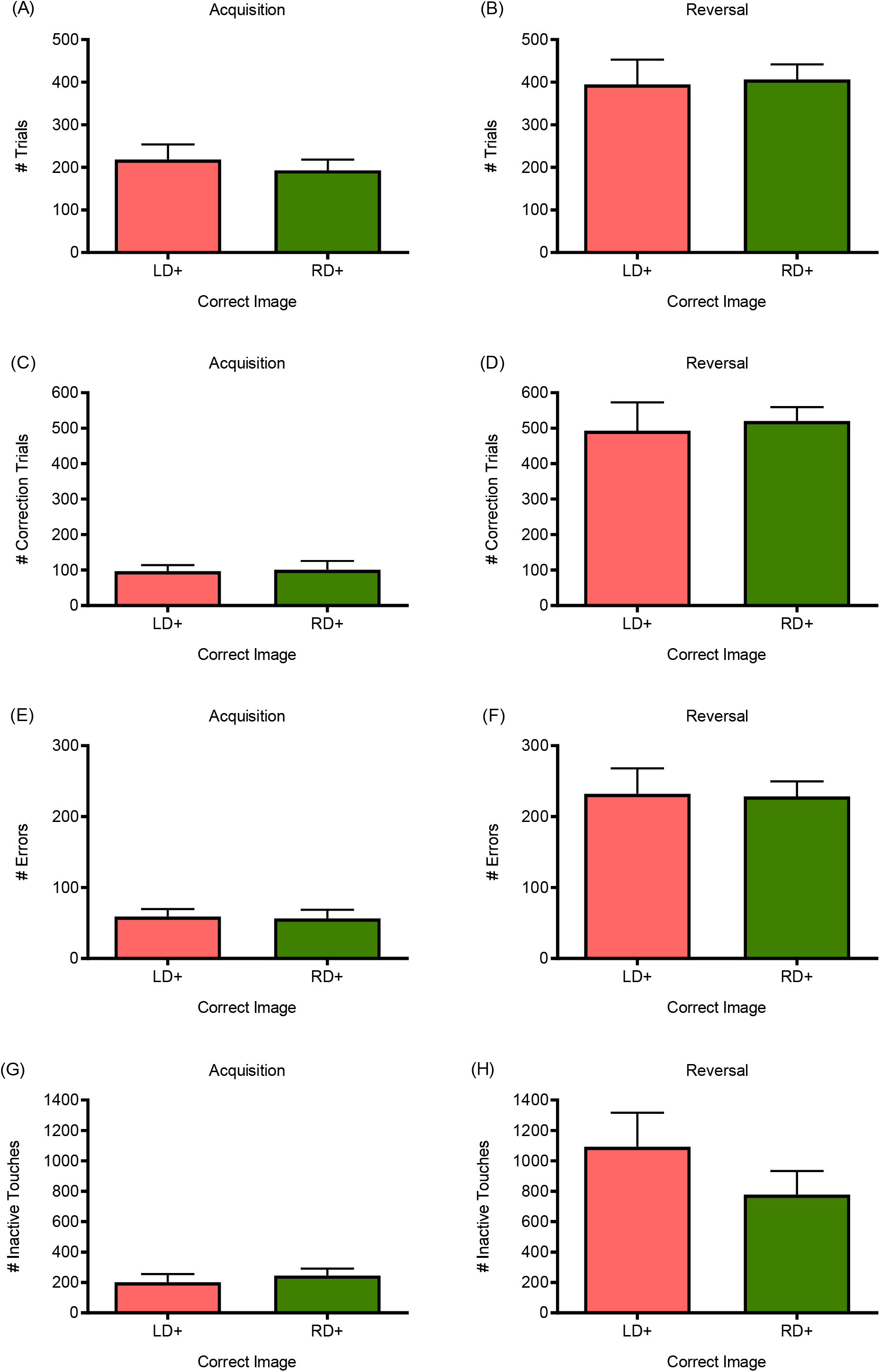
Both groups in the mirrored diagonal bar image pairing perform similarly by all accessory measures examined. As with the spider-plane image pairing, both LD+ (pink bars) and RD+ (green bars) groups in the mirrored diagonal pairing complete the same number of trials overall in both acquisition (A) and reversal (B). However, unlike the spider-plane image pair, there were no significant differences between LD+ and RD+ for any of the other examined measures, including total number of correction trials required during acquisition (C) and reversal (D), total number of errors made in acquisition (E) and reversal (F), and total inactive touches in acquisition (G) and reversal (H). Data are presented as mean ± SEM

Of note is that across all these measures, the diagonal groups in acquisition had fewer correction trials, errors, and inactive touches than either the diagonal groups in reversal or the spider and plane groups in both acquisition and reversal. However, this is likely an effect of the mice in the mirrored diagonal groups being over-trained on the touchscreen task. To minimize animal use in this study that was peripheral to our group’s initial experimental purpose (probing for cognitive defects), these mice were the same as those used in the mirrored plane pairing after a two-day break. Thus, though these animals had never acquired a paired stimuli-reward rule, they were more familiar with the touchscreen experimental apparatus than mice usually are when beginning acquisition.

### 3.6 Spider image stimulus is twice as bright as the opposing plane stimulus

To explore possible reasons for the bias we observed using the standard plane-spider stimuli pairing, we next examined the relative luminance of the touchscreen images used throughout our studies. Given they are all binary images (images where each pixel is either black or white), the number of white pixels is an accurate indicator of luminosity. While both the plane and spider images were the same size at 240×240 pixels, the spider stimulus has nearly twice as many white pixels as the plane (13,414 vs 7,776). As expected, comparison of the mirrored diagonal images (220×220 pixels) showed the exact same number of white pixels for both stimuli at 26,036, a notably higher number than that of both the plane and spider images.

## 4 DISCUSSION

The pairwise visual discrimination test is one of the most basic in the Bussey-Saksida system test battery. It effectively assesses rodent recognition memory and cognitive flexibility (reversal phase). The animal needs to choose between two images juxtaposed on the screen, nose-touching the correct image to earn a reward, usually a small amount of palatable food or liquid. Selecting the incorrect image results in a brief timeout. The most commonly used image pairs are spider-plane ^11,12,19,22^ and marble-fan ^25,36-39^. Our data presented here call attention to an apparent image bias in the spider-plane pairing, which could pose a problem for accurate data analysis.

Our study here shows that when the spider of the spider-plane pair is rewarded, mice reach criterion faster in both the acquisition and reversal phases, at least in our mouse line of interest. Interestingly, this pattern could also be seen in preliminary data from three other touchscreen pairwise discrimination studies at our facility, which was revealed in a simple meta-analysis of the anonymized data **(Table 1)**. While most of these were small pilot studies that were not continued after initially seeing no genotypic differences (and thus underpowered to reliably detect statistical significance here), analyzing the data by rewarded image shows that spider-correct group required fewer days to reach criterion than the plane-correct group in all three lines in both acquisition and reversal. These data indicate the presence of a strong bias in the spider-plane image pairing that can’t be attributed to the use of a specific mouse line. Still, even though our study was powered enough to reach significance, the data we present strongly suggests the need for more comprehensive studies into more diverse and widely relevant mouse lines by groups dedicated to touchscreen cognitive paradigms.

**Table 1.** Days to Criterion

| Anonymized<br>Mouse Line<br>ID | Acquisition,<br>Plane+<br>(Days) | Acquisition,<br>Spider+<br>(Days) | Reversal,<br>Plane+<br>(Days) | Reversal,<br>Spider+<br>(Days) |
| --- | --- | --- | --- | --- |
| A | 11.73 | 10.00 | 15.20 | 14.13 |
| B | 12.71 | 10.66 | 16.33 | 12.57 |
| C | 13.50 | 9.83 | 13.50 | 11.75 |

Another important measure in the reversal phase of this task is days to 50%. This is the number of days it takes the animal to return to baseline and begin performing at chance level. When the learned rule is initially switched on the mice in the reversal phase, they perform well below chance, indicating that they are perseverating on the old rule. These “perseveration errors” are distinct from so-called “learning errors” (errors made when an animal is learning a new paired association) and have been shown to be mediated by distinct brain regions. The days to 50%, then, becomes a good indicator of cognitive flexibility, or the ability of the animal to abandon the previously learned rule and adapt to a new one, a related but distinct cognitive process. ^40,41^. Our data show that the spider-correct group in the reversal phase require significantly fewer days to 50% than the plane-correct group. These data suggest that image bias not only affects measures of learning and memory but may also confound the analysis of cognitive flexibility and obscure the discovery of subtle cognitive phenotype abnormalities in test animals.

The above data are also a good indication that the normal approach of counter balancing may not be sufficient to counteract bias in all studies. Counter balancing is a method by which the rewarded “correct” image is always randomized among test subjects in such a way that half the animals start with one rewarded image choice in the acquisition phase (e.g. spider), while the other half starts with the opposite image choice as the rewarded one (e.g. plane). This is a commonly used strategy to eliminate the argument that any image bias is the driving factor behind reported data. However, our results here suggest this may not be enough when there is a dominant bias. It could also make the data more difficult to interpret. For example, days to 50% won’t just be a measure of cognitive flexibility if there is also an effect of image bias driving the amount of time it takes for the mouse to switch rules. The fact that our spider-correct group reached this reversal milestone faster than the plane-correct group suggest this is the case.

We also show that the bias is detectable as early as the first day of the acquisition phase, a fact that presents more problems for accurate data interpretation. In a normal learning curve for acquisition, mice should start around chance performance (50% correct) and progress to passing scores (above 80% correct) after numerous days of training. Several mice in the spider-correct group failed to show this typical learning curve, some of which were already performing above 80% correct on the first day, as indicated in the individual learning curves in the supplementary data. On the other hand, no mice in the plane-correct group showed such anomalous learning behavior, a fact that is further reflected in the averaged learning curves for the two groups presented above.

In addition to days to criterion, image bias also affects number of correction trials and errors made, both of which were significantly lower for the spider-correct group during acquisition. The number of inactive touches, or how often the mouse interacts with the screen when no images are displayed, also tended to be lower for the spider-correct group in acquisition and reversal. This is an especially striking effect of correct image group considering that this is a measure of behavior in the absence of image presentation. It suggests the image bias effect is confounding elements of the experiment unrelated to direct stimulus presentation. Even when no image is projected on the screen, the two groups show significantly different behaviors in approaching the task. These data imply that a strong image bias may not only directly affect overall task performance as measured by days to criterion but also how the animals approach and attend to the task in the moment. This presents another important dimension to the task that may not be adequately controlled for with the standard technique of counterbalancing stimuli as discussed above. Our review of the touchscreen literature was unable to find an in-depth analysis of such peripheral measures and their interpretations, suggesting another component of such tasks that stands to benefit greatly from more close analysis by those developing these tools for more widespread use by casual users such as the authors of the present study.

Further complicating the issue of bias is that as far as we can find, only one early touchscreen study published an analysis of bias, examining multiple image pairings^18^. In this study, one of six image pairings analyzed was reported to have a bias and this pairing was eliminated in the study moving forward. However, the approach in this study suffered from a few procedural limitations. For one, the study was done in rats, and to our knowledge this was never followed up in mice, who may have quite different visual capacities. For another, the researchers in this study set up arbitrary thresholds as indicated by a significant deviation from chance (50%), and they only reported a bias when one image in the image pairing was selected more often than the top threshold (around 65%) and the opposing image was selected less often than the lower threshold (around 35%). If these thresholds weren’t reached, the pairing was considered unbiased even if the data present in such a way that a bias seems evident if a standard statistical comparison (such as a Student’s T-test) had been used between images in the image pairing. Finally, the researchers chose to evaluate bias with only one testing session, rather than a series of them over a number of days, as the discrimination task is normally done. Thus, they fail to see differences influenced by a bias that might become more apparent over multiple testing sessions, as seen by the more dramatic differences seen in our averaged learning curves and Kaplan-Meier curves.

Without the luxury of testing large numbers of image pairs like these ourselves, our solution to eliminate image bias was simply using two of the same images turned at different angles. This was seen to be effective in both our mirrored-plane and mirrored-diagonal bars experiments. And while we observed that in the instance of the mirrored-planes, some image pairings can lack enough distinguishing features for the mice to effectively learn the paired association rule, our mirrored-diagonal patterns showed that with enough salient features in the lower half of the image screen (a region that is potentially more accessible to the mouse field of view), the mice can reach criterion when non-geometric images are used. Importantly, there were no significant differences between the left diagonal bar correct and right diagonal bar correct groups by any of the measures that we saw differences in the spider-plane pairing. These include days to criterion, days to 50% correct (reversal), the averaged learning curves, Kaplan-Meier curves, total number of corrections trials, total errors, and total inactive touches. This image pairing thus provides a clear solution to eliminating the bias seen in the more dissimilar standard pairings such as spider and plane. And while we are not the first to use these diagonal bar patterns, to our knowledge we are the first to report it as an effective control for bias seen in non-geometric shape image pairings such as spider and plane.

But what is the cause of the spider-plane image bias we see? An investigation into potential explanations provides evidence that it may in part be a product of luminosity differences between the spider and plane stimuli. As we report above, the spider image has nearly twice the number of white pixels as the plane image despite both image files being the same size. This suggests the experimental animals may have been drawn toward the brighter image. This could be for a few reasons. One is that it may be a more salient stimuli and thus draw more attention at baseline. Another possibility, one more specific to the pairwise discrimination task, is that it may be because during pre-training, when the mice are familiarized with the touchscreen apparatus before acquisition, the mice learn to interact with the screen by being trained to ignore the blank screen and touch the screen lit by an image to receive a reward. In fact, as indicated in Methods, the mice spend the last four of the five pretraining stages discriminating between a blank screen and one with a randomized pretraining image: a brighter screen and a darker screen. And while this is an important step in familiarizing the mouse with utilizing the touchscreen, it is possible that when the mouse moves on to the acquisition phase and must choose between the spider and the plane, they are following an unintended association of “brighter screen is the rewarded stimulus,” thus showing a tendency to approach the spider image more often. Regardless, the mirrored diagonal bar pairing had identical white pixel densities, as expected for reflected images, eliminating any brightness effect. They also had a much higher white pixel density than either the plane or the spider, perhaps making the images a more engaging and salient stimuli pairing. This, in turn, may facilitate discrimination between the two despite their marked similarity, something that was not possible for the animals in the mirrored plane experiment. Of course, this is just an interpretation based on our necessarily limited study. A more complete answer will require ongoing study by those with more resources and expertise in the touchscreen cognition field, as well as those studying rodent visual perception.

As already mentioned, we are not the first to use such simple geometric shapes or striped patterns. Similar striped images have been used elsewhere ^28,35^. Other simple images have also been used in the visual discrimination task and include + and x ^34^ and “x” and “=“, possibly due to the obvious easiness to matched the shapes for illumination ^42^. The use of simple images, such as stripes, X/=, and +/x, have the advantages of being both equiluminant and equally complex, minimizing the chance that the mice are relying on extra visual cues that could be prone to bias. However, not much is known about their effectiveness in more complicated tasks. For example, most studies on paired association learning (dPAL) utilized flower, spider, and plane ^24,26,43-51^, and we were only able to find one study that used stripes for the dPAL task ^52^. It is a pure speculation that complex images may be beneficial for complex tasks such as PAL and transitive inference. However, this is an important question for future studies to address. In these more complicated tasks, an image bias in even just one pairing could cause a domino effect that greatly muddles results. For example, given the design of the dPAL task, two out of six additional trial types could be affected if there is an image preference that encourages approaching the spider more readily than the plane.

Clarifying these experimental difficulties, however, will rely not only on accurate image selection, but also on a better understanding of rodent visual perception. Considering the important role of visual ability in learning and memory in mice, particularly in tasks such as these, the findings that visual abilities differ among strains ^53^ and decline with age ^54^, and the intriguing yet under-appreciated research on species differences in visual perception in mice and humans ^55-57^, much more research is needed to ensure that visual stimuli used in mouse behavioral tests are species-appropriate and do not introduce salient confounding factors. For example, the touchscreen method has been reported to be effective in testing rodents thought to have poor enough vision to prohibit learning visual cognitive tasks ^18^. But if these visually impaired animals are learning associations based on differences in overall brightness levels rather than finer image features, the task and images used should be more tightly controlled to take this into account. Along those lines, many touchscreen studies do not report images used. Until more is known about rodent visual perception to enable completely bias-proof image selection, our limited analysis suggests that it might be necessary for authors to specify images in their publications.

Furthermore, the visual features that create a salient bias are likely different across species, complicating the translational potential of touchscreen results when a bias exists. While humans use a complex assortment of visual features such as contrast, shape, brightness, size, and more to distinguish images, animal models such as mice may primarily rely upon only one or a subset of these. Two images that seem different, yet unbiased enough for a discrimination task by human observers might still suffer a bias in mice if not perfectly matched for a visual feature (like brightness) that may be the primary property rodents rely on for image discrimination. These translational differences are vital considerations when comparing a primarily olfactory animal like a mouse with a primarily visual animal like a human. Taking this into account, it seems especially important to ensure perfect equivalence in as many features as possible to guarantee that the results obtained in a visual cognitive task such as touchscreen pairwise discrimination are primarily *cognitive* and not *visual*.

While we are not the first to warn against the possibility of bias or observe an image bias in pairwise discrimination testing using the touchscreen method, to our knowledge we are the first to analyze the phenomenon more deeply to uncover the numerous measures the bias is capable of confounding, at least in one mouse line. And while other researchers have used these mirrored images in previous studies, to our knowledge we are the first to propose them as a specific solution to the problem of image bias. One important thing to note in closing is that it is unlikely a confounding image bias, such as the one we report here, is responsible for any positive results that have been seen in previously published touchscreen studies. Of greater concern is that a strong image bias such as that we found with the spider-plane pairing could introduce enough noise into touchscreen studies of cognition to decrease the task’s sensitivity in detecting more subtle cognitive differences, resulting in the observation of false negatives. Given that higher level cognition is already so difficult to investigate in rodents in a translationally meaningful way, it is vital that such useful and increasingly used paradigms as the touchscreen platform are optimized to make the task as sensitive as possible. Eliminating confounding image biases in oft-used image pairings and consistently updating these based on advances in understanding rodent visual perception are important steps in this direction. Doing so will require the effort and expertise of both touchscreen cognitive study pioneers and rodent visual perception researchers to take noted observations like those presented here and study them with more comprehensive experiments on image bias.

## Notes

### Competing Interest Statement

The authors have declared no competing interest.

